# Decline in pneumococcal disease in unimmunized adults is associated with vaccine-associated protection against colonization in toddlers and preschool-aged children

**DOI:** 10.1101/293266

**Authors:** Daniel M. Weinberger, Virginia E. Pitzer, Gili Regev-Yochay, Noga Givon-Lavi, Ron Dagan

## Abstract

Vaccinating children with pneumococcal conjugate vaccines disrupts transmission, reducing disease rates in unvaccinated adults. When considering changes in vaccination strategies (e.g., removing doses), it is critical to understand which groups of children contribute most to transmission. We used data from Israel to evaluate how the build-up of vaccine-associated immunity in children was associated with declines in IPD due to vaccine-targeted serotypes in unimmunized adults. Data on vaccine uptake and prevalence of colonization with PCV-targeted serotypes were obtained from a unique study conducted among children visiting an emergency department in southern Israel and from surveys of colonization from central Israel. Data on invasive pneumococcal disease in adults were obtained from a nationwide surveillance study. We compared the trajectory of decline of IPD due to PCV-targeted serotypes in adults with the trajectory of decline of colonization prevalence and trajectory of increase in vaccine-derived protection against pneumococcal carriage among different age groupings of children. The declines in IPD in adults were most closely associated with the declines in colonization and increased vaccination coverage in children in the range of 36-59 months of age. This suggests that preschool-aged children, rather than infants, are responsible for maintaining the indirect benefits of PCVs.

---

Pneumococcal conjugate vaccines (PCVs) have had a well-documented impact on the incidence of invasive pneumococcal disease (IPD) and pneumonia in young children (1–3). PCVs reduce the burden of disease in two ways: they directly protect vaccinated individuals who are exposed to the bacteria against invasive infections, and they indirectly protect vaccinated and unvaccinated individuals (including adults) by reducing the prevalence of carriage of vaccine-targeted serotypes and thus reducing transmission. Colonization of the nasopharynx of young children represents the main reservoir for transmission of pneumococcus, and PCVs reduce the proportion of children who are colonized with serotypes targeted by the vaccine (4). The reduction in the burden of disease in unvaccinated adult age groups resulting from this indirect protection greatly outweighs the reduction in the burden of disease seen in vaccinated children alone (5).
While PCV programs have effectively reduced the burden of disease in many countries, the cost of the vaccine remains a major concern in both affluent and resource-poor settings. This issue has taken on particular urgency as some lower-income countries “graduate” from being eligible for financial support from Gavi.

To reduce costs while maintaining widespread use of the vaccine, there has been interest in reducing the number of doses of PCVs delivered to children (6–8). Such a strategy could be used in populations that have already achieved strong reductions in disease due to the vaccine. Currently, most countries use either a 3+0 schedule (3 doses in the first six months of life, no booster dose) or a 2+1 schedule (2 doses in first six months of life, 1 booster dose administered at 9-15 months of age). The newly proposed schedule would include just one primary dose and one booster dose (1+1). This schedule is being considered in the United Kingdom and is being evaluated by the World Health Organization. Reduced-dose schedules have been shown to be immunogenic (7, 9, 10). However, because of the importance of indirect protection that results from the use of PCVs, it would be desirable for any new dosing strategy to be able to maintain indirect protection (6).

To consider this issue, it is critical to determine which groups of young children contribute most to the indirect benefit of the vaccine for unimmunized adults. Epidemiological and modeling studies of transmission focused on households and daycare centers suggest that toddlers and older children, rather than infants, drive transmission in the population (11–14). Likewise, the decline in IPD due to vaccine-targeted serotypes in adults is delayed and slower than the decline observed in vaccinated children (15). This might indicate that the transmission benefit (i.e. indirect effect) of vaccinating children during the first year of life is not realized until a few years later when those children reach an older age group.

In this study, we evaluated how the build-up of vaccine-associated immunity against colonization in different age categories was associated with declines in IPD due to vaccine-targeted serotypes in unimmunized age groups. We used a unique and ongoing survey of children in Israel that allowed us to quantify vaccine uptake and IPD rates over time to evaluate these associations.

## METHODS

### Data sources

PCV7 was introduced into the national immunization program in Israel for all children in July 2009 with a catch-up campaign for all children <24 months of age and was gradually replaced with PCV13 starting in November 2010. Carriage and vaccine uptake data for children were obtained from an ongoing study of children visiting the emergency department at Soroka University Medical Center (16). Each weekday, the first four Jewish children and first four Bedouin children under the age of 5 years who visited the emergency department (ED) were enrolled in the study. A nasopharyngeal swab was collected and cultured, and serotype was determined using Quellung reactions, as described previously (16). For each child enrolled in the study, the number of doses of PCV7 and PCV13 received was recorded. These vaccine data were previously used to represent PCV uptake in Israel, since data from the region are within the range of the average vaccine uptake nationwide (17). We calculated uptake of PCV7/13 in each month post-PCV introduction in age bands that varied in width and ages included. Data on IPD in adults were collected as part of a national surveillance system in Israel (18).

For our primary analyses, only data from Jewish individuals were included due to different demographics of the minority populations in southern Israel (where the carriage data were drawn from) compared with the entire country (where the IPD data were drawn from). As a further evaluation of changes in prevalence among healthy children (rather than among children visiting the ED), we evaluated changes in the prevalence of PCV7-targeted serotypes among healthy children living in central Israel who were sampled as part of a series of cross-sectional surveys of nasopharyngeal colonization (19, 20). The study was approved by the Soroka University Medical Center Ethics Committee and the Sheba Medical Center Ethics Committee.

### Calculation of the “Population Direct Effect” against colonization

The observed impacts of vaccination depend upon both the direct and indirect effects of the vaccine (21). To determine which age groups are key to determining the overall impact of vaccination, it is necessary to differentiate between the direct and indirect protection among children of different ages. In practice, interpreting the association between direct protection against carriage in different groups of children and IPD patterns in adults can be confounded by transmission between groups of children (**Figure S1**). For instance, if the vaccine predominantly disrupts carriage in toddlers, and toddlers are the main source of transmission to both infants and the elderly, it can appear that declines in carriage in infants are the drivers of the declines in the elderly (**Figure S1**). However, carriage in infants in that schematic would simply be an intermediate step on the casual pathway between vaccination of toddlers and declines in disease in infants.

To avoid this issue of confounding by indirect protection, we can calculate the ‘population direct effect,’ which provides an estimate for the overall effect of the vaccine on carriage that would be expected in the absence of indirect protection (22). The quantity is simply a function of the individual-level direct efficacy of the vaccine (as measured in a randomized controlled trial) and the proportion of the population that is vaccinated. The population direct effect would typically be calculated by estimating the proportion of individuals in a particular age strata that received 1, 2, or 3+ doses of vaccine and multiplying this by the individual-level vaccine efficacy of 1, 2, or 3 doses against colonization due to vaccine-targeted serotypes. We modified this calculation to allow for waning of vaccine-derived protection. For a given age band (*a*) and time point (*t*):

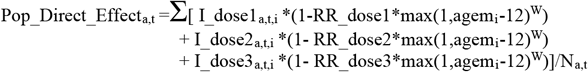

This quantity is summed across all individuals in a particular age band surveyed during a specific month. I_doseK_a,t,I_ is an indicator for whether an individual had received 1, 2, or 3+ doses of PCV7/13; agemi is the age in months of the individual; ‘W’ is the rate of waning and is set to 0.11 per month based on a meta-analysis of waning protection against carriage (23); and N_a,t_ is the number of individuals surveyed in a particular month in a particular age band. We assume no waning of vaccine efficacy during the first year of life, followed by protection that wanes at a constant rate after 12 months of age, regardless of number of doses they have received. Estimates of vaccine efficacy (1-RR_doseK) were obtained from a randomized controlled trial of PCV7 among children in Israel that used a variety of dosing schedules (13, 24). The point estimate for efficacy against colonization in the first year of life was estimated at 0%, 27%, and 46% for 1, 2, or 3 doses, respectively. To account for effects of catch-up vaccination, we made some simplifying assumptions based on previous studies (13): For children who had received 1 dose and who were over the age of 12 months, we assumed they had initial efficacy equal to an infant who had received 2 doses (27%). For children who received 2 doses and who were over the age of 12 months, we assumed these were both catch up doses, with protection equal to a child who had completed the full 3-dose series (46% initial efficacy). In sensitivity analyses, we varied the rate of waning and the estimated efficacy, but these did not qualitatively alter the conclusions of the main analysis.

The observed estimates of the population direct effect for any given time point and stratum were based on small numbers, so we used cubic splines to smooth the trajectory of the population direct effect. This was accomplished using PROC GAM in SAS v9.4 (Cary, NC), where the outcome was the observed estimates of the population direct effect, and time was modeled with a cubic spline with 3 degrees of freedom. Separate smoothing models were fit for each age range.

### Evaluating the association between direct protection and indirect effects

We fit Poisson regression models where the outcome was the number of cases of IPD due to PCV7-targeted serotypes in a particular month and specific adult age group, and the sole covariate was the Pop_Direct_Effect_a,t_, corresponding to a specific age band <5 years old. We controlled for seasonality using monthly dummy variables. We only used data from the Jewish population for this analysis. We compared the likelihood of each model given the data based on the Akaike Information Criterion (AIC) (25). Model likelihoods were calculated by comparing the AIC score for model *i* with the AIC score from the best-fitting (lowest AIC score) model (25):

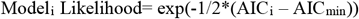

By convention, models that were within 2 AIC points of the best model were considered to not be meaningfully different from the best model (26). To obtain a summary measure of the importance of each age group (i.e., 0-5m, 6-11m, 12-17m), we averaged the likelihoods for all models in which the age band for the population direct effect variable covered that 6-month age group.

### Evaluating the association between carriage in different groups of children and IPD in adults

The relationship between carriage prevalence in children and IPD in adults can be difficult to interpret due to indirect protection (**Figure S1**). Nonetheless, carriage prevalence provides the most directly observable measure of vaccine-associated changes in transmission in children. We fit Poisson regression models where the outcome was the number of IPD cases due to PCV7-targeted serotypes in a particular month and adult age group. The sole covariate was log(carriage prevalence) in a given age band at each time point. Log(carriage prevalence) was smoothed using PROC GAM, as described above. For these analyses, we focused on PCV7 serotypes only (rather than the six additional serotypes that are were added into PCV13). This is because there was a brief period of time between the introduction of PCV7 and PCV13, which made it difficult to disentangle early serotype replacement from vaccine-associated effects. Since the PCV7 serotypes are present in both vaccines, focusing on these serotypes allowed for more interpretable trajectories.

## RESULTS

### Description of the population

Nasopharyngeal swabs and vaccine status were obtained from 4,464 Jewish children <5 years of age visiting the ED between 2009 and 2016. Among these children, the prevalence of PCV7 serotypes declined from 21% in 2009 to 2.6% in 2016. These declines were apparent in all age groups of children <5 years of age, with the most rapid initial declines among 12-23 and 24-35 month old children and slower declines among <12 month and 36-59 month olds (**Figure 1, S2**). The differences in these trajectories between age groups were similar in healthy Jewish children living in central Israel (**Figure S3**). During the same time period, the proportion of IPD cases in adults caused by vaccine-targeted serotypes declined (**Figure 2**). These declines coincided with a rapid increase in vaccine uptake among children.

**Figure 1:**
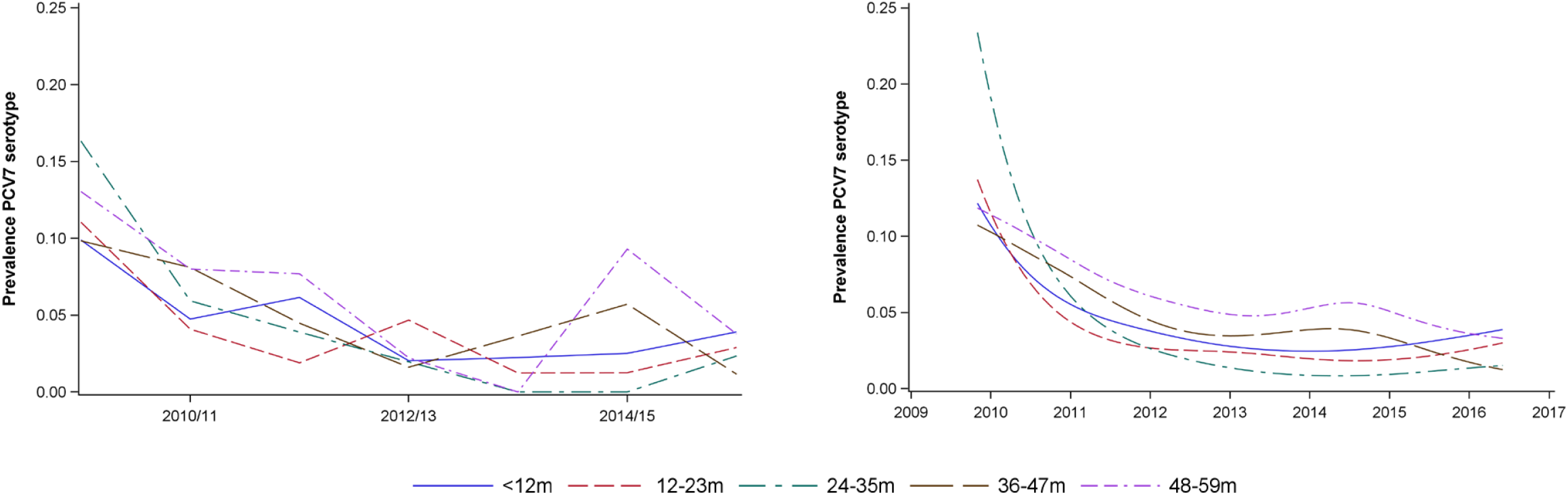
(A) Observed proportion of nasopharyngeal swabs that were positive for PCV7 serotypes by epidemiological year (July-June) among Jewish children <5 years of age, stratified into 1-year age categories (<12m, 12-23m, 24-35m, 36-47m, 48-59m). (B) Smoothed proportion of children carrying PCV7 serotypes in each month 2009-2016.

**Figure 2:**
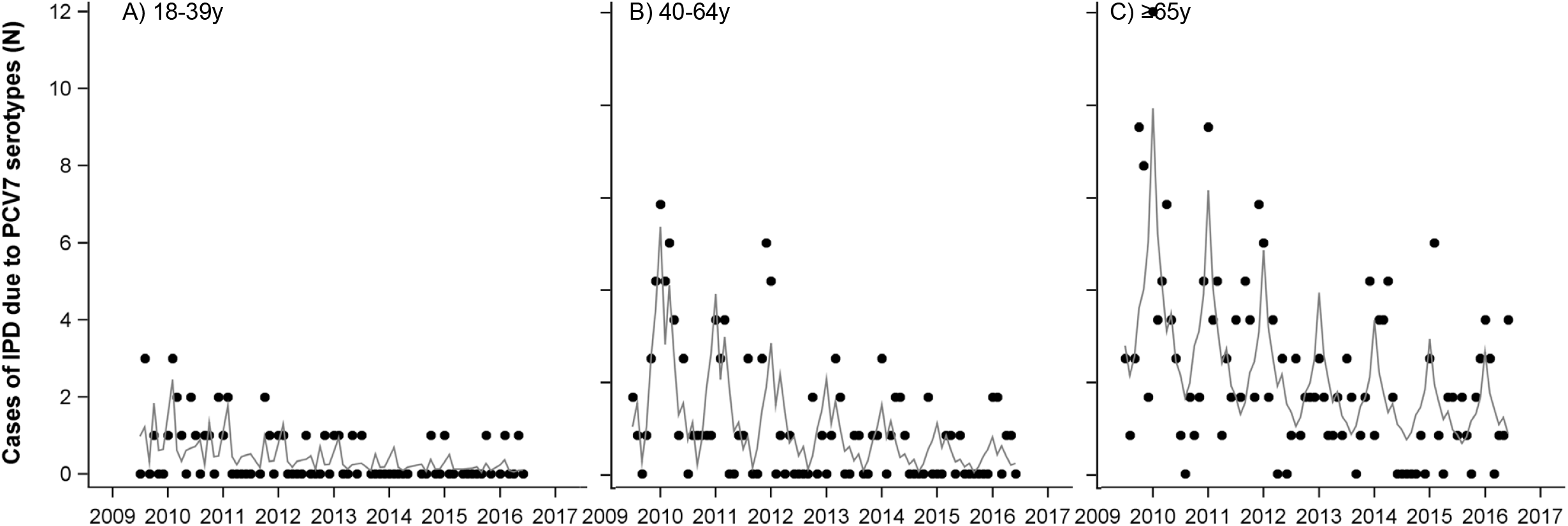
Number of cases of invasive pneumococcal disease due to PCV7 serotypes among Jewish adults in Israel, July 2009-June 2016. The observed number of cases in each age group is indicated by the dots. A smoothed trend (fit with a generalized additive model with a spline for time and monthly dummy variables) is shown with the gray line.

### Estimated increase in population direct protection against carriage by age group

We estimated the reduction in the prevalence of PCV7-targeted serotypes that would be expected in the absence of an effect of the vaccine on transmission (the “population direct effect”). Among children <12 months of age, the population direct effect reached a plateau at the beginning of the study period and remained stable at 15-20% (**Figure 3**). This is because the vaccine effectiveness against carriage of the two primary doses received in the first year of life is low, and uptake of these two doses among infants was stable throughout the study period. Among 12-23 month old children, the population direct effect against carriage increased rapidly following vaccine introduction and plateaued at around 30% by 2012. Among older age groups, the increase in the population direct effect was delayed until cohorts of children vaccinated as infants aged into the specified age band (**Figure 3**). The maximum population direct effect achieved was lower in older age groups due to waning of vaccine-induced immunity.

**Figure 3:**
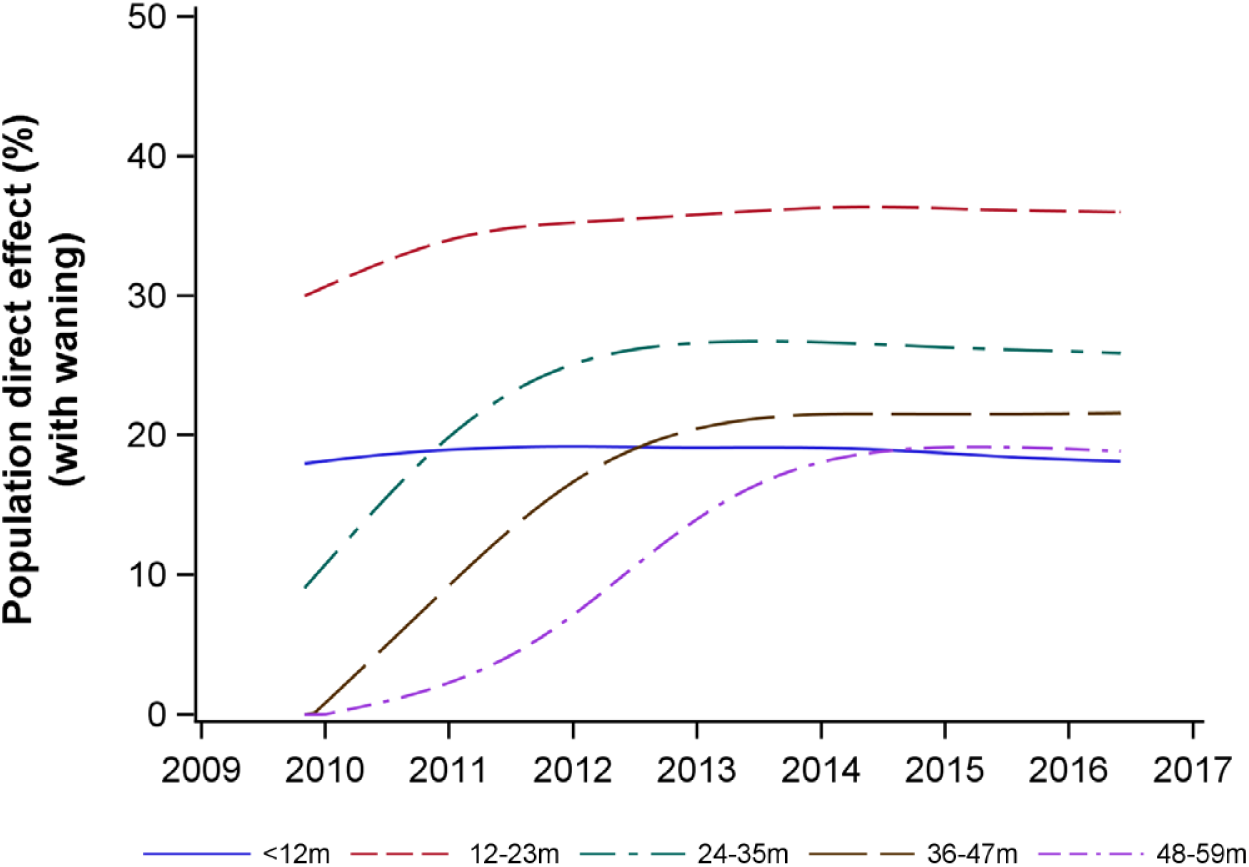
Increase in the population direct effect against carriage by age group for Jewish children: <12m, 12-23m, 24-35m, 36-47m, 48-59m. The population direct effect represents the expected vaccine effectiveness against carriage if there was no effect of the vaccine on transmission. Estimates were smoothed with splines prior to plotting

### Increase in direct protection against carriage is associated with indirect protection against IPD

We evaluated the association between the decline in the occurrence of IPD in adults and the population direct effect against carriage in different age bands of children <5 years of age. In **Figure 4**, each horizontal band indicates an age range in which the population direct effect was calculated. Horizontal bands that are higher on the y-axis indicate a better fit to the decline in IPD due to PCV7 serotypes in adults. Bands highlighted in color and above the dashed line had strong statistical support (≤2 difference in AIC score from the best-fitting model). This demonstrates that the decline in IPD due to PCV7 serotypes among adults 18-39 y, 40-64 y and ≥65 y of age was most strongly associated with the increase in the population direct effect among 36-54 month, 30-59 month, 30-54 month old children, respectively. However, several age bands covering the range from 6-59 months of age had strong statistical support. It is notable that when the age bands included children <12 months old, the association was significantly weaker. When looking in aggregate across all combinations of age bands (**Figure 4B**), there was strong evidence that the direct protection among older children (i.e. ≥36 months) was important to explaining the decline in IPD among adults, as well as strong evidence that direct protection among children <12 months was not as important; the role of direct protection among 12-35 month old children was ambiguous.

**Figure 4:**
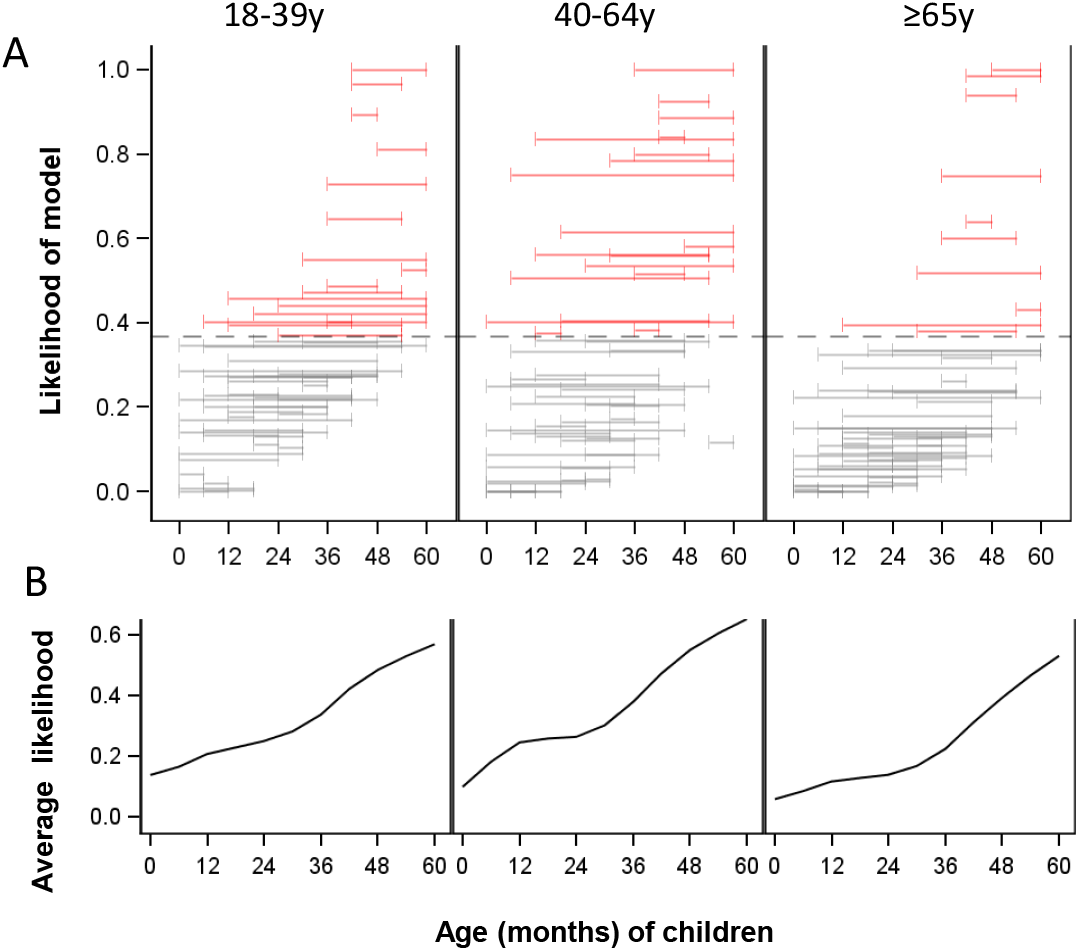
Relative goodness of fit between the population direct effect in different age ranges of children (indicated by the horizontal bars) and invasive pneumococcal disease (IPD) in adults. (A) Each horizontal bar indicates a specific age range in which the population direct effect was calculated based on uptake of 1, 2, and 3 doses of the vaccine in that age group and expected efficacy of these doses against colonization. The vertical position of the bar along the y-axis indicates the goodness of fit, as measured by the likelihood of the model given the data compared to the best-fitting model (which has a relative likelihood of 1). These values are calculated from the Akaike Information Criteria score. The population direct effect in age ranges that are placed higher on the y-axis fit the adult IPD data better. Bars colored in red were not meaningfully different from the best-fit model (AIC score within 2 points). (B) The average goodness of fit at each age.

### Declines in carriage in infants and older children are associated with declines in disease in adults

We expect that increasing vaccine-induced (direct) protection causes declines in IPD in adults by reducing carriage in children. We therefore also evaluated the association between the decline in IPD due to PCV7-targeted serotypes in adults and the carriage prevalence of PCV7-targeted serotypes at each time point in different age groups of children. Declines in carriage of PCV7-targeted serotypes in older children (≥24 months) and children <6 months of age were most strongly associated with the declines in IPD due to PCV7-targeted serotypes in adults (**Figure 5**). However, there was variability between the 18-39y, 40-64y, and ≥65y age groups in which age bands of children were most strongly correlated. These patterns could be due to the direct or indirect protection of these age groups (**Figure S1**) and should therefore be interpreted with caution. Directly linking carriage prevalence with the population direct effect, the decline in carriage prevalence in children <6 months of age was most strongly correlated with increasing population direct protection in children ≥12 months old (**Figure S4**).

**Figure 5:**
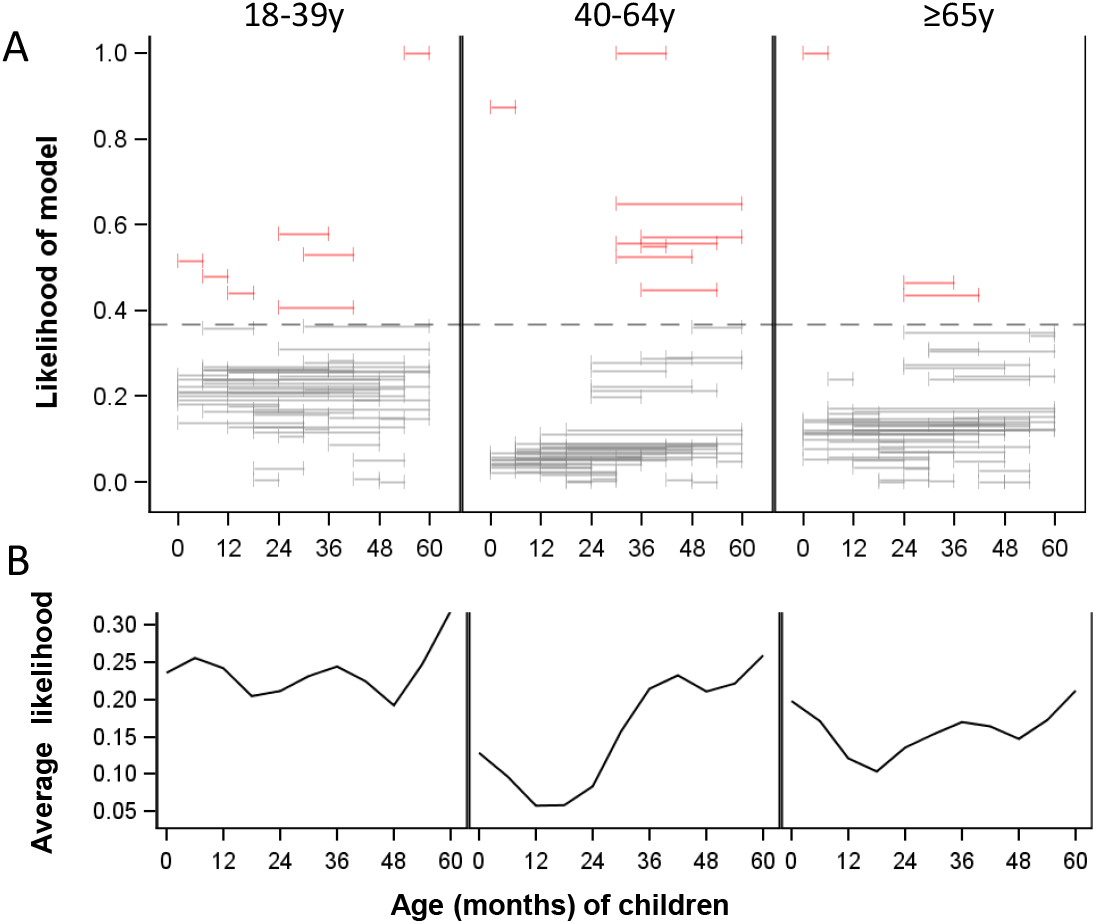
Relative goodness of fit between prevalence of PCV7 serotypes among healthy children in different age ranges (indicated by the horizontal bars) and invasive pneumococcal disease (IPD) in adults. (A) Each horizontal bar indicates an age range in which (smoothed) carriage prevalence was calculated. The vertical position of the bar along the y-axis indicates the goodness of fit, as measured by the likelihood of the model given the data compared to the best-fitting model (which has a relative likelihood of 1). These values are calculated from the Akaike Information Criteria scores. Carriage prevalence in age ranges that are placed higher on the y-axis fit the adult IPD data better. Bars colored in red were not meaningfully different from the best-fit model (AIC score within 2 points (B) The average goodness of fit at each age.

## DISCUSSION

Indirect protection has been a critical component of the overall impact of PCVs, as evidenced by the large declines in IPD among older adults who were not recipients of the vaccine. The patterns of association described here between vaccine-induced direct protection, carriage prevalence in children, and IPD patterns in adults suggest a mechanism for indirect protection. In particular, the analysis of population direct effects suggests that increasing vaccine-induced protection in toddlers and pre-school-aged children drives down carriage prevalence in these older children and among children <6 months of age. These declines in carriage among children then drive declines in IPD in adults. Therefore, direct protection of toddlers and preschool-aged children is the main causal driver of declines of IPD in adults, and carriage in infants <6 months old is either an intermediate step on the causal pathway or due to confounding, but is probably not the main causal driver (**Figure S1**). Such a mechanism is consistent with the delays in indirect protection observed in adults in several high-income populations following the introduction of PCV (15, 27).

These findings have important implications for the possible success of reduced-dose vaccination schedules. Our findings suggest that vaccine uptake in infants plays a minor role in influencing the indirect effects of PCVs. This is partially because the two primary doses of the vaccine received in the first year of life have a weak effect on colonization. Rather, it is the third dose that provides strong protection against carriage and subsequent transmission. This suggests that if the booster dose in a 1+1 schedule provides adequate protection against colonization, and uptake rates of the vaccine in toddlers and older children are sufficiently high, then the reduced dose schedule should be effective at maintaining indirect protection in populations resembling the one in our study. Recent immunological data suggests that the 1+1 and 2+1 schedules provide comparable levels of immunity after the booster dose, as measured by serum IgG (7). If these serum IgG levels correlate with mucosal immunity in the nasopharynx, the reduced-dose schedule would be expected to provide effective protection against colonization that is comparable to a 2+1 schedule in these older children. These findings also suggest that catch-up campaigns targeting 1-5 year old children could help to accelerate the realization of the indirect benefits of PCVs. Further analyses of the timing of declines in IPD from settings that did or did not use catch-up vaccination could help to further evaluate this issue.

Our results are consistent with previous epidemiological and modelling studies of pneumococcal colonization that suggest that toddlers and young school-aged children, rather than infants, drive transmission of pneumococcus in the population (11–14). A study of the impact of PCV7 in the United States demonstrated strong declines in IPD among <12 month and 12-23 month old children immediately after introduction of the vaccine in the 2000 calendar year and declines in 24-35 month old children, 40-64 year old adults, and ≥65 year old adults in the 2001 calendar year (27). Likewise, previous analyses demonstrated that the decline in IPD in adults in the US was delayed by ~1 year compared to the decline in IPD in children <5 years old (15). These patterns reinforce the notion that it is important to maintain strong immunity to colonization in these older children in order to maintain indirect protection of younger children and adults.

Our analyses focused primarily on the Jewish population in Israel, which has a population structure and sociodemographic profile similar to that of populations in high-income countries in Europe and North America. The contact structure, age-specific prevalence, and intensity of transmission will be different in a low-income setting. As a result, the age groups that contribute most to transmission and indirect protection could differ in low-income settings as well. Performing analyses of the dynamics of timing of initiation of indirect effects in lower-income populations and in countries with or without catch-up campaigns could help to further evaluate the issue of which age groups contribute most to indirect protection.

A strength of this study is the ability to extract data on vaccine uptake and colonization at a high resolution of time and age from a large sample of children. Thus, we were able to monitor the increases in uptake, changes in carriage prevalence, and declines in vaccine-targeted serotypes at an unusually detailed level across the population. Furthermore, the IPD data were obtained from a robust national surveillance system, allowing us to quantify the indirect effects of the vaccine in adults. Limitations of the data include our reliance in the main analysis on carriage and vaccine uptake data from a study performed in an emergency department setting. However, our secondary analysis of carriage data from children living in central Israel (**Figure S3**) suggests that the trajectory of decline of PCV-targeted serotypes is similar in the general pediatric population. Finally, we have used a relatively simple set of analyses based on regression models to evaluate this issue. Transmission patterns are complex and dynamic, and an age-structured transmission model could be used to further evaluate the importance of these different age cohorts in different populations and to evaluate the potential impact of changes to the vaccine schedule.

In summary, our findings suggest that direct protection of toddlers and preschool-aged children with PCVs are most influential in maintaining indirect protection against IPD among older adults in Israel. Any changes in vaccine schedules should focus on maintaining immunity in these older children.

---

**Figure S1:**
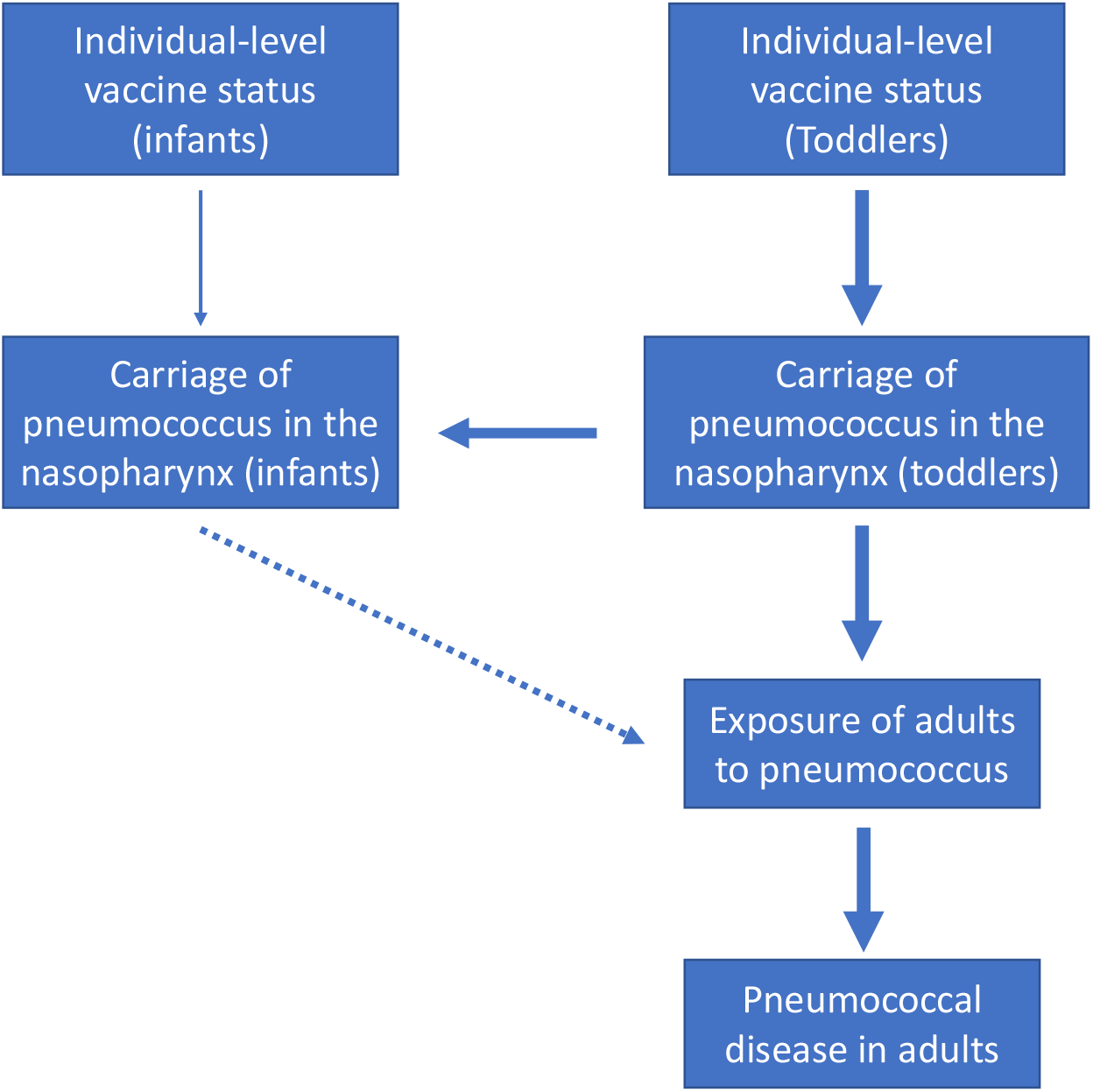
Schematic causal diagram illustrating the effect of vaccination of different age groups on direct and indirect protection.

**Figure S2:**
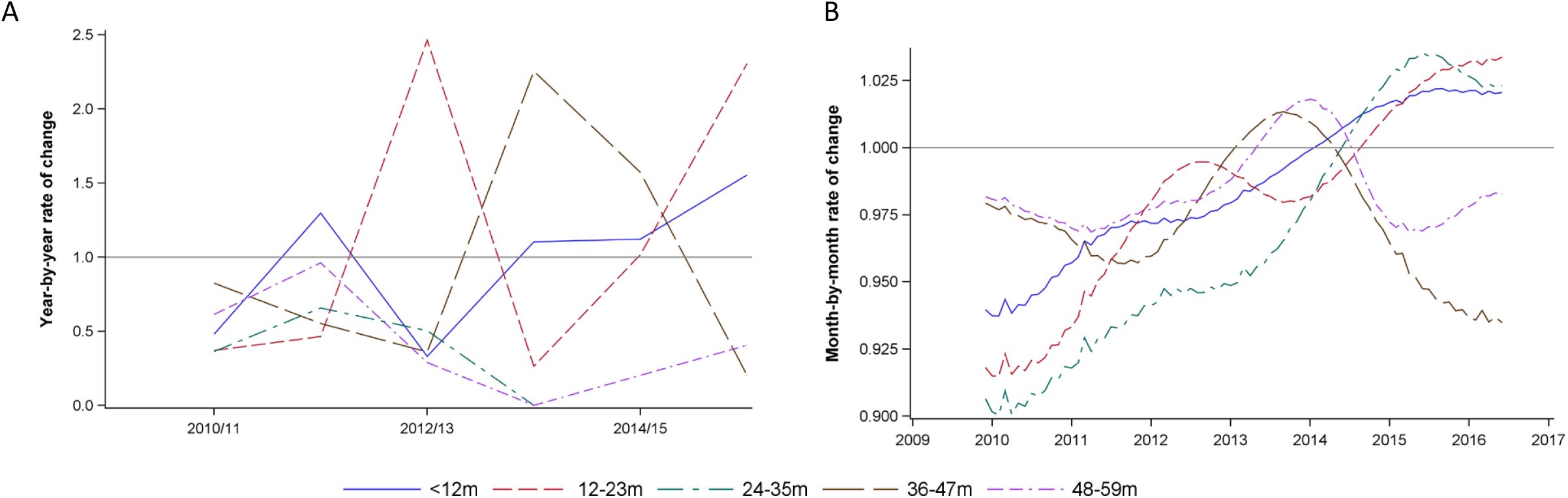
Rate of change of carriage prevalence, corresponding to those seen in Figure 1. Values further below 1 indicate faster declines. (A) The ratio of prevalence in in the indicated epidemiological year compare with the previous epidemiological year, using observed data. (B) The ratio of prevalence in the indicated month vs the previous month, using smoothed prevalence curves.

**Figure S3:**
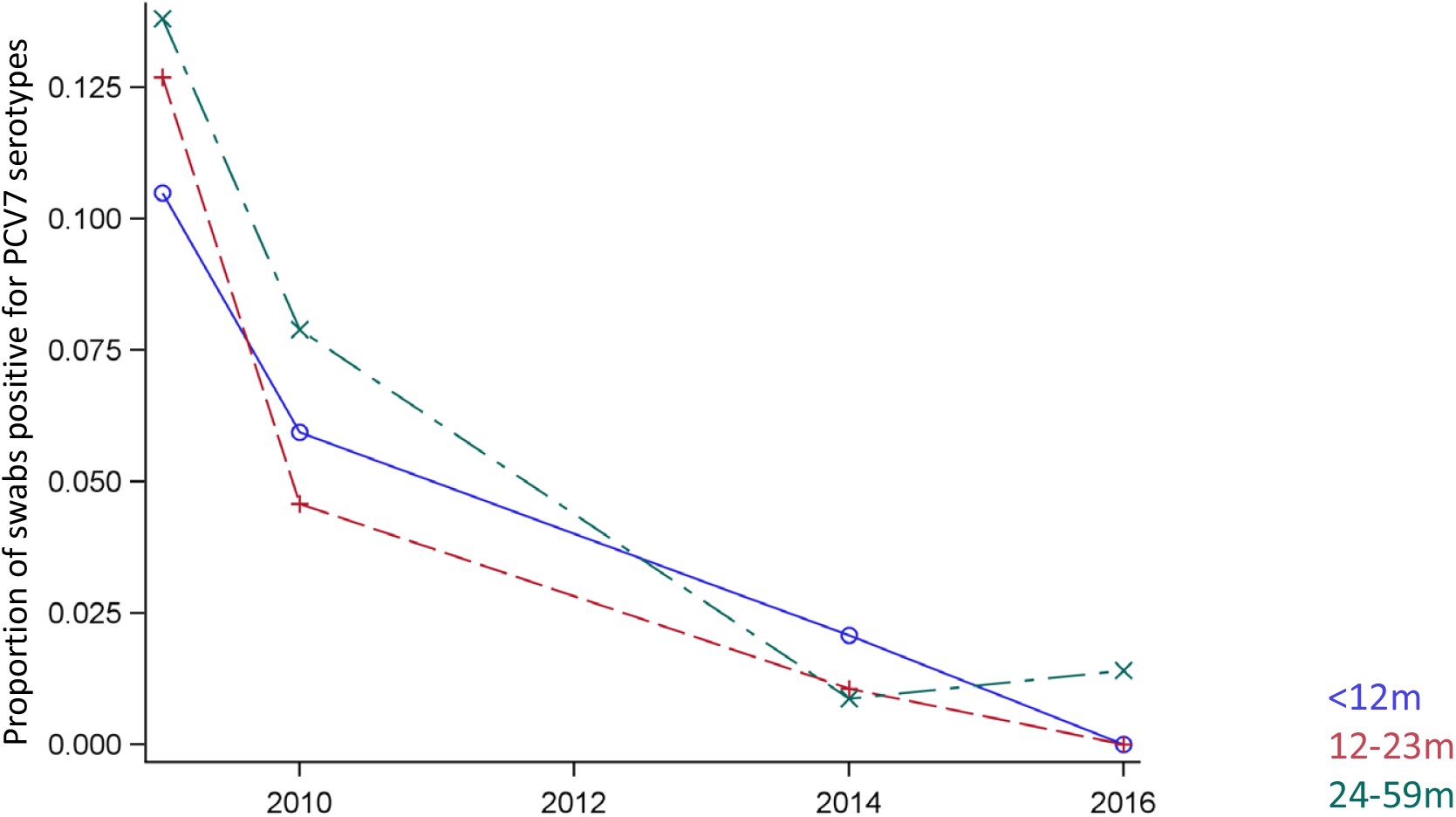
Observed proportion of nasopharyngeal swabs that were positive for PCV7 serotypes by calendar year among Jewish children <5 years of age living in central Israel, stratified into age categories (<12m, 12-23m, 24-59m).

**Figure S4:**
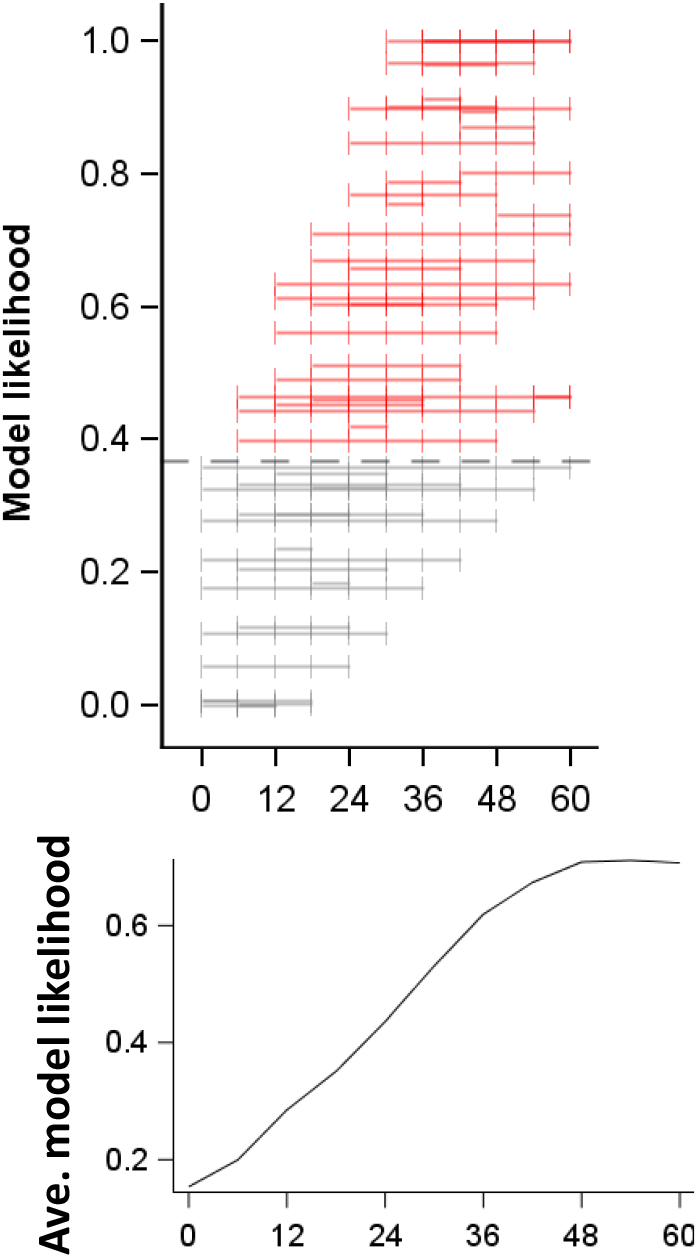
Relative goodness of fit between the population direct effect in different age ranges of children (indicated by the horizontal bars) and the prevalence of PCV7 serotypes among children <6 months of age. Each horizontal bar indicates an age range in which the population direct effect was calculated. The vertical position of the bar along the y-axis indicates the goodness of fit, as measured by the likelihood of the model given the data compared to the best-fitting model (which has a relative likelihood of 1). These values are calculated from the Akaike Information Criteria scores. Population direct effect estimates for age ranges that are placed higher on the y-axis fit the adult IPD data better. Bars colored in red were not meaningfully different than the best-fit model (AIC score within 2 points). Some bars had near-equivalent livelihoods and overlapping ranges, each age range only has hash marks at the end of the bar. The bottom panel represents the average goodness of fit at each age.

## Acknowledgements

2) All authors have seen and approve of the manuscript. DMW conceived of and performed the analyses. DMW, VEP designed the analyses. RD, NGL, GRY collected the data. DMW wrote the first draft of the paper. All authors contributed to revisions of the paper

3) Research reported in this manuscript was supported by the National Institute of Allergy and Infectious Diseases of the National Institutes of Health (NIH/NIAID) under award number R01AI123208 and by the Bill and Melinda Gates Foundation (OPP1176267). The content is solely the responsibility of the authors and does not necessarily represent the official views of the National Institutes of Health. VEP acknowledges support from NIH/NIAID (R01AI112970) and the Bill and Melinda Gates Foundation (OPP1116967). GRY acknowledges support from the NIHP (25–10). The studies in Israel from which the databases from southern Israel were derived were partially funded by Pfizer (Grant no. 0887X1-4603)

4) The authors thank Dr. Gail Rodgers and Ms. Prachi Vora for discussion of the figures and results.

6) Preliminary results from this study were presented at the International Symposium for Pneumococci and Pneumococcal Diseases, Melbourne Australia, April 2018.

7) Conflicts of Interest: Within the last two years, DMW has received consulting fees from Affinivax and Pfizer. RD has received grants and consulting and speaker fees from Pfizer; grant and consulting fees from MSD and consulting fees from MeMed. GRY has received consulting fees from Pfizer and research support from Pfizer and MSD.

